# Effects of woody riparian vegetation on macroinvertebrates are context-specific and large in urban and especially agricultural landscapes

**DOI:** 10.1101/2022.06.22.497242

**Authors:** Martin Palt, Daniel Hering, Jochem Kail

**Affiliations:** Department of Aquatic Ecology, University of Duisburg-Essen, 45117 Essen, Germany; Environmental Campus Birkenfeld, University of Applied Sciences Trier, 55761 Birkenfeld, Germany; Centre of Water and Environmental Research, University of Duisburg-Essen, 45117 Essen, Germany

**Keywords:** agricultural landuse, urbanisation, macroinvertebrates, river restoration, woody riparian vegetation

## Abstract

1. Woody riparian vegetation (WRV) benefits benthic macroinvertebrates. However, in recent large scale studies, the effects of WRV on macroinvertebrates were small compared to catchment landuse, raising the question about the relevance of WRV in restoration. Limited effects of WRV might be due to context specificity: While some functions are provided by WRV irrespective of catchment landuse, others depend on the landscape setting.
2. Recursive partitioning modelling was used to identify context dependent effects of WRV on streams macroinvertebrates’ ecological status in small lowland (n = 361) and mountain streams (n = 748). WRV cover was quantified from orthophotos along the near (500 m) and far (5,000 m) upstream river network and used to predict the site’s ecological status. Agricultural, urban and woodland cover at the local and catchment scales along with hydromorphology were considered as partitioning variables.
3. In rural agricultural landscapes, the effect of WRV on the ecological status was large, indicating that establishing WRV can improve the ecological status by as much as two classes.
4. In streams impacted by catchment urbanization, effects of WRV were largest, but WRV cover and ecological status were both low, indicating practical limitations of WRV restoration in urban catchments.
5. *Synthesis and applications:* Independent effects of WRV on macroinvertebrates’ ecological status can be discerned from catchment landuse. While WRV can also improve the ecological status in urban settings, it is especially relevant for river management in rural agricultural catchments, where developing WRV potentially are effective measures to reach good ecological status

## 1. Introduction

Woody riparian vegetation (WRV) benefits aquatic ecosystem health in temperate regions, where most streams and rivers are naturally bordered by trees (Ellenberg, 1988). This notion is supported by a large number of studies demonstrating functional links between WRV and ecosystem processes in the riparian and aquatic environment (reviewed e.g. in Broadmeadow & Nisbet, 2004; Sweeney & Newbold, 2014).

Several of these functions are provided irrespective of landscape settings while others are context-specific and linked to adjacent landuse in the floodplain. Independently from adjacent landuse, WRV provides organic material like leaves, twigs, and large wood that serve as food and habitat for different aquatic organisms (Oelbermann & Gordon, 2000) and redirect flow, creating higher flow- and substrate-diversity and channel features like pools, bars and undercut banks (McBride et al., 2010). Moreover, herbaceous bank vegetation is supressed, promoting natural channel patterns and dynamics (Parkyn et al., 2005). Finally, WRV serves as habitat and as migration or dispersal corridor for terrestrial invertebrates, birds, mammals and terrestrial life-stages of aquatic insects (e.g. Petersen et al., 2004; Van Looy et al., 2014). In principle, these functions depend on the presence of trees alone. In contrast, retention of nutrients, fine sediments, and pesticides is also related to inputs from adjacent agricultural areas and strongly increases with WRV width (Arora et al., 2010; Gericke et al., 2020; Ramesh et al., 2021). Moreover, some functions of WRV are more relevant in specific contexts. For example, shading limits primary production and reduces water temperature, which is especially relevant if elevated nutrient levels would otherwise result in excessive phytobenthos and macrophyte growth (Kiffney et al., 2003; Nebgen et al., 2019). Besides landscape setting, these effects also depend on the length of WRV patches. While shading by WRV causes lower equilibrium water temperatures within few hundred meters (Kail et al., 2021), the positive effect of reducing inputs of nutrients, fine sediment and pesticides rather accumulates over long distances. Since several functions depend on landscape setting and length of the WRV patches, these should be considered when investigating the effect of WRV on river biota. Based on these reasons, higher effects are expected in agricultural catchments, as well as from wider and longer WRV sections along the riparian corridor.

A large number of reach-scale empirical studies have found positive effects of WRV on functional traits and community composition of benthic macroinvertebrates while limited effects were evident in some recent, larger-scale empirical studies. In reaches bordered by WRV, shares of shredding macroinvertebrates were higher than in open reaches (ZumBerge et al., 2003; Thomas et al., 2016; Turunen et al., 2019), indicating the role of WRV to provide leaves as a food source (Lecerf & Richards, 2010). Biomass and abundance of macroinvertebrates were lower in shaded stream reaches due to lower water temperature and light availability (Smith, 1980; Noel et al., 1986; Kaylor & Warren, 2018), limiting instream primary production (Parkyn et al., 2003; Feld et al., 2011). With decreasing canopy cover, the abundance of tolerant taxa like Chironomidae and Oligochaeta strongly increased on the expense of sensitive taxa like Plecoptera and Ephemeroptera (Kiffney et al., 2003; Thompson & Parkinson, 2011). The decline of sensitive taxa also reflects the associated increase in fine sediment input and substrate siltation (Davies & Nelson, 1994). Additionally, sensitive taxa benefit especially from the retention of pesticides by WRV (Bunzel et al., 2014).

These reach-scale studies usually compare differing configurations of WRV and often follow a BA/CI design. This implies that larger-scale stressors originating from the catchment are similar. However, recent studies, which analysed a larger number of reaches from different catchments, indicate that catchment landuse as a proxy for larger-scale stressors superimposes on the effects of riparian landuse cover on benthic macroinvertebrate traits (Le Gall et al., 2021; Palt et al., 2022). Therefore, catchment landuse must be considered when investigating the effect of WRV on benthic macroinvertebrates. From a management perspective, these studies imply that establishment of WRV would not substantially raise the ecological status if catchment landuse remains unchanged.

Most of the studies mentioned above investigated the effect of reach scale WRV on functional traits and community composition, yet there is limited knowledge of the effect on the ecological status according to the EU’s Water Framework Directive (LeGall et al., 2022; Palt et al., 2022; Tolkkinen et al., 2021). The few studies using comparable indices of macroinvertebrate communities’ naturalness like the Index of Biotic Integrity (IBI) or a Quantitative Macroinvertebrate Community Index (QMCI) found however better conditions in reaches with WRV compared to those lacking it (e.g. Newbold et al., 1980; Parkyn et al., 2003; ZumBerge et al., 2003; Aschontis et al., 2016).

Yet, besides aforementioned large-scale stressors superimposing on the effects of riparian landuse, also the low effectiveness of reach-scale restoration is often attributed to stressors acting at the catchment scale (Jähnig et al., 2010). This raises the question, under which conditions WRV, a widely used restoration measure, can significantly improve the ecological status of macroinvertebrate communities. Most studies on the effect of reach scale WRV on macroinvertebrates quantified landuse on low-resolution data, covering forested areas but not including small patches of WRV like single lines of trees along rivers (Dahm et al., 2013; Lorenz and Feld, 2013; Tolkkinen et al. 2021). Moreover, riparian corridors investigated in these studies were wide (50 – 100 m), rather reflecting forest cover in the whole floodplain and adjacent hillslopes. Many functions like shading mainly depend on WRV directly adjacent to the river banks (Kail et al., 2021) and wide strips of WRV can hardly be established in densely populated regions or areas intensively used for agriculture. Therefore, studies are missing that include small woody patches and focus on WRV effects in a narrow riparian corridor which is important from an ecological and management point of view.

Against this background, this study aims at identifying conditions, under which WRV in a narrow riparian corridor has significant effects on the ecological status of macroinvertebrates using high resolution data on WRV. We hypothesise that the effect of WRV is context-specific and differs with catchment and local landuse, length of the considered riparian corridor and hydromorphology. More specifically, we expect WRV having its largest effects in agricultural landscapes, because several of its functions described above are mainly linked to agricultural landuse in the floodplain. WRV even far upstream is expected being important, since positive effects of some functions potentially accumulate downstream. Meanwhile, stressors related to urban catchment landuse like point source pollution and stormwater runoff are expected to limit effects of WRV.

## 2. Material and Methods

### 2.1. Biological data

Data on macroinvertebrate samples from small lowland (n = 361; 18–189 m MSL) and small mountain streams (n = 748; 58–594 m MSL), taken between 2004 and 2013, were acquired from three German federal state: Hesse, North Rhine-Westphalia, and Saxony-Anhalt (Fig. 1). Sites in lowlands and mountains were analysed separately due to assumed differences in the interaction of aquatic and terrestrial environments based on factors such as topography, discharge, slope and consequently flow velocity and stream morphology.

**Fig. 1:**
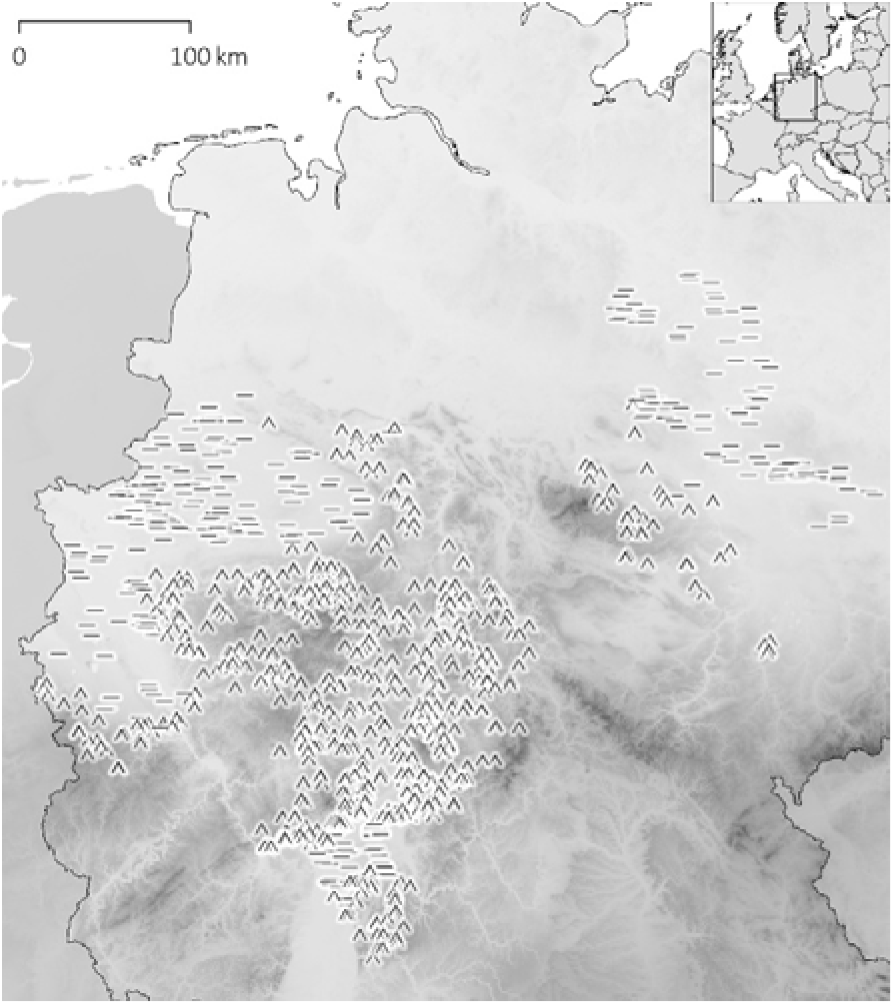
Location of macroinvertebrate sampling sites in Germany: Bars show lowland and chevrons mountain streams.

Macroinvertebrate samples were taken according to the multi-habitat sampling method described in Haase et al. (2004). The species-level taxa lists were processed using the online tool PERLODES (https://www.gewaesser-bewertung-berechnung.de/index.php/perlodes-online.html), which amongst others computes the river-type specific multimetric index (MMI). The MMI is the core component of the ecological status assessment according to EU Water Framework Directive in Germany. The MMI reflects the impact of various stressors like hydromorphological degradation, altered hydrology and impacts of landuse (Böhmer et al., 2004).

The dataset was pre-processed to exclude data of insufficient quality: Only samples with at least 5 taxa and samples taken between December 1^st^ and April 30^th^ were included to guarantee reliability and comparability. For the same reason samples with a saprobic index > 2.7 were excluded, as theses correspond to polluted streams affected by point sources. Sites with barriers within 5,000 m upstream of the sampling site were excluded, since these trap sediments, alter the thermal regime to varying degrees, and therefore potentially mask sediment retention and shading by WRV.

### 2.2. Riparian landuse

Upstream riparian buffers were demarcated for each sampling site at two spatial scales, starting at the sampling site and extending for 500 m and 5,000 m upstream length, respectively (Fig. 2), referred to as near upstream and far upstream in the following. Riparian buffers were delineated using ESRI ArcView (Version 3.3) and included tributaries. Laterally, they covered 30 m to either side starting from the stream banks, hence excluded the water surface, quantifying terrestrial landuse only. Water surfaces were taken from official ATKIS landcover data (www.adv-online.de/Products/Geotopography/ATKIS). For small streams not included as water surfaces in ATKIS, the wetted width was approximated by a mean width measured from orthophotos for all different Strahler orders (n = 30 each). The rather small buffer width of 30 m was chosen, as it is relevant in river management and restoration and because many functions like shading mainly depend on woody riparian vegetation (WRV) directly adjacent to the stream.

**Fig. 2:**
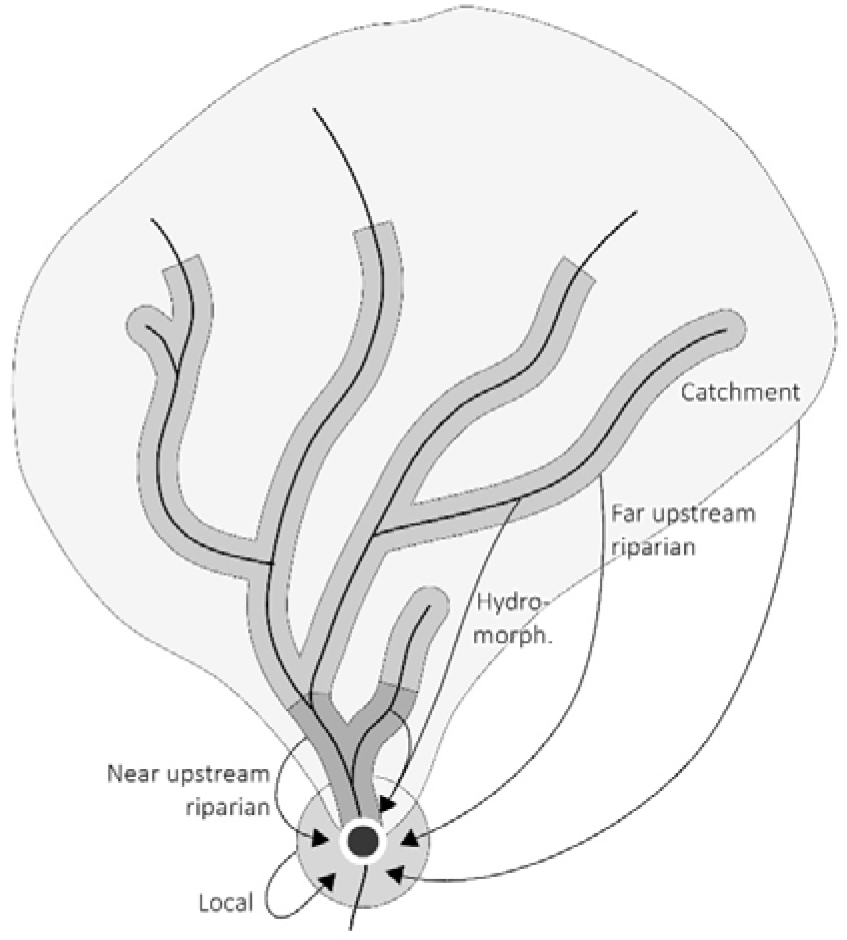
Landuse was assessed in the catchment, locally in a 250 m circle around the sampling site and at two lengths in the riparian corridor upstream from the sampling site 30 m to either side of the stream: Near upstream extents for 500 m, and far upstream for 5,000 m respectively. Available hydromorphological assessments were aggregated for both upstream lengths. Not to scale.

WRV was quantified from ATKIS data. Its detailed landuse classes were grouped into seven categories: (1) “arable land”, (2) ‘‘grassland”, (3) ‘‘natural vegetation”, (4) “urban green space”, (5) “urban”, (6) “water surface”, and (7) “woody vegetation”, with some rare landuse classes excluded (e.g. quarries, harbours). Given the minimum size of woody vegetation patches in ATKIS is 0.1 ha, smaller landscape features, like single lines of trees along rivers, were missing. Therefore, ATKIS data in the riparian corridor were complemented by WRV, down to single trees, identified on orthoimages. These were obtained from the German Federal Agency for Cartography and Geodesy and were mostly CIR and some RGB images with a 0.2 m resolution (0.4 m for some few older RGB images). Only orthoimages taken between April and August and closest to 2010 were used to match the vegetation period and macroinvertebrate samples respectively.

Orthoimages were processed in an object-based image analysis (OBIA), consisting of image segmentation and classification of resulting objects. The multiresolution segmentation into objects of homogenous pixel patches was carried out in Trimble’s eCognition (Version 9.3.0) based on the pixel values of the colour bands. For their classification a support vector machine (SVM) classifier was developed based on a training dataset of 40 representative orthophotos (n = 14 RGB, n = 26 CIR), which had been classified in a supervised semi-manual nearest neighbour classification approach. The SVM classifier distinguished woody vegetation, other forms of vegetation (grassland, cropland), and non-vegetated areas (built-up areas or bare soil) based on shape, colour and brightness of the objects, as well as the Visible-band Difference Vegetation Index (VDVI, RGB images) or Normalized Difference Vegetation Index (NDVI, CIR images). This SVM classifier was applied to the orthophotos using the R package e1071 (version 1.7-3). General accuracy of segmentation and classification was assessed visually. Additionally, accuracy of the SVM classifier was assessed using cross-validation on the training dataset. Woody vegetation objects identified on the orthoimages replaced ATKIS landuse patches of the categories, “arable land”, “grassland”, “natural vegetation”, “urban green space”, and “urban”. Improving the spatial resolution of landuse data in close proximity to the river was a prerequisite to correctly quantifying the percentage cover of near and far upstream WRV.

### 2.3. Catchment and local landuse

For each sampling site, landuse outside the riparian corridor was quantified at two spatial scales (Fig. 2). (1) The catchment, i.e. drainage basin to the sampling site, was delineated on a digital elevation model (DEM, 10 m resolution) and visually checked. (2) The local surroundings of the sampling site were a circular buffer with a radius of 250 m.

Percentage cover of the three landuse categories “urban”, “agriculture”, and “woodland” was quantified for each scale with ESRI’s ArcGIS Desktop 10.8. Urban landuse comprises all built-up areas and infrastructure. It has detrimental effects on stream biota from catchment (e.g. impervious surfaces) to local scale (e.g. light pollution). Agricultural areas are subject to tillage, fertilization, and pesticide application, which respectively may result in inputs of fine sediments, nutrients, and toxic substances. Woodlands are the predominant potential natural vegetation in temperate regions and should cause the least detrimental effects approximating natural instream conditions. Quantifying woodland cover at catchment and local scale allows distinguishing the effect of woody riparian vegetation in the riparian buffer from adjacent woodland cover, i.e. forest cover in general.

### 2.4. Hydromorphology

Stream morphology pre-sets the potential of transport of sediment or detritus, as well as the thermal regime. Therefore, the effect of woody riparian vegetation (WRV) might further depend on instream hydromorphology.

Hydromorphology mappings and assessment results following Gellert et al. (2014) were provided by regional authorities. Twenty-five individual hydromorphological parameters are mapped for 100 m river segments and compared to natural reference conditions. Their deviation from reference conditions is assessed on an ordinal scale ranging from unchanged with just minor deviations (class 1) to heavily degraded (class 7). Scores of the 25 parameters are aggregated to main parameters: (1) “channel pattern”, (2) “longitudinal profile”, (3) “channel bed features”, (4) “cross section”, (5) “channel bank features”, and (6) “floodplain conditions”. For each sampling site, mean assessment scores for main parameters 1 to 5 were aggregated based on all available assessment segments 500 m and 5,000 m upstream of the sampling sites (Fig. 2). Main parameter “floodplain conditions” was omitted not to duplicate information on riparian vegetation.

### 2.5. Statistical analysis

Model-based recursive partitioning (Zeileis et al., 2008) was used to test the hypotheses. Its core was a linear regression model (*1m*) for the macroinvertebrate multimetric index (MMI) given the percentage cover of woody riparian vegetation (WRV) in the near upstream and far upstream riparian buffer, which is fitted per maximum likelihood estimation. The other variables in the data set, namely urban, agricultural and woodland cover at both the local and catchment scale, as well as the hydromorphological assessment results at the near and far upstream scale, were incorporated as candidate partitioning variables.

The recursive approach first tests for the entire dataset if the estimates of the *1m* show any significant parameter instability towards the gradients of any candidate partitioning variable. If statistically significant instability is found (Andrews’ *supLM* test; Zeileis, 2005), the optimal split in the gradient of the partitioning variable causing the highest parameter instability is calculated. This split point optimizes the maximum likelihood for the core model fitted to the resulting child datasets. The process is reiterated until no more parameter instability with respect to the candidate partitioning variables in the *1m* is found for the thus final subdatasets.

The recursive splitting of the entire dataset can be intuitively displayed in a tree-diagram similar to other CART approaches. However, this method differs from most of these as it does not partition the data into groups of observations with similar response values. Rather it splits the data into groups of observations with similar model trends between the response (MMI) and core predictors (WRV) not used for partition (Garge et al., 2013).

Spearman’s *ρ* correlation coefficient between WRV and woodland cover in the catchment were calculated in order to assess if potential effects of WRV on the MMI were independent or rather a proxy for effects of larger-scale forest cover.

## 3. Results

### 3.1. Lowland streams

The lowland sampling sites were split into three subdatasets (LL.1 – LL.3) by recursive partitioning based on two partitioning variables (Fig. 3).

**Fig. 3:**
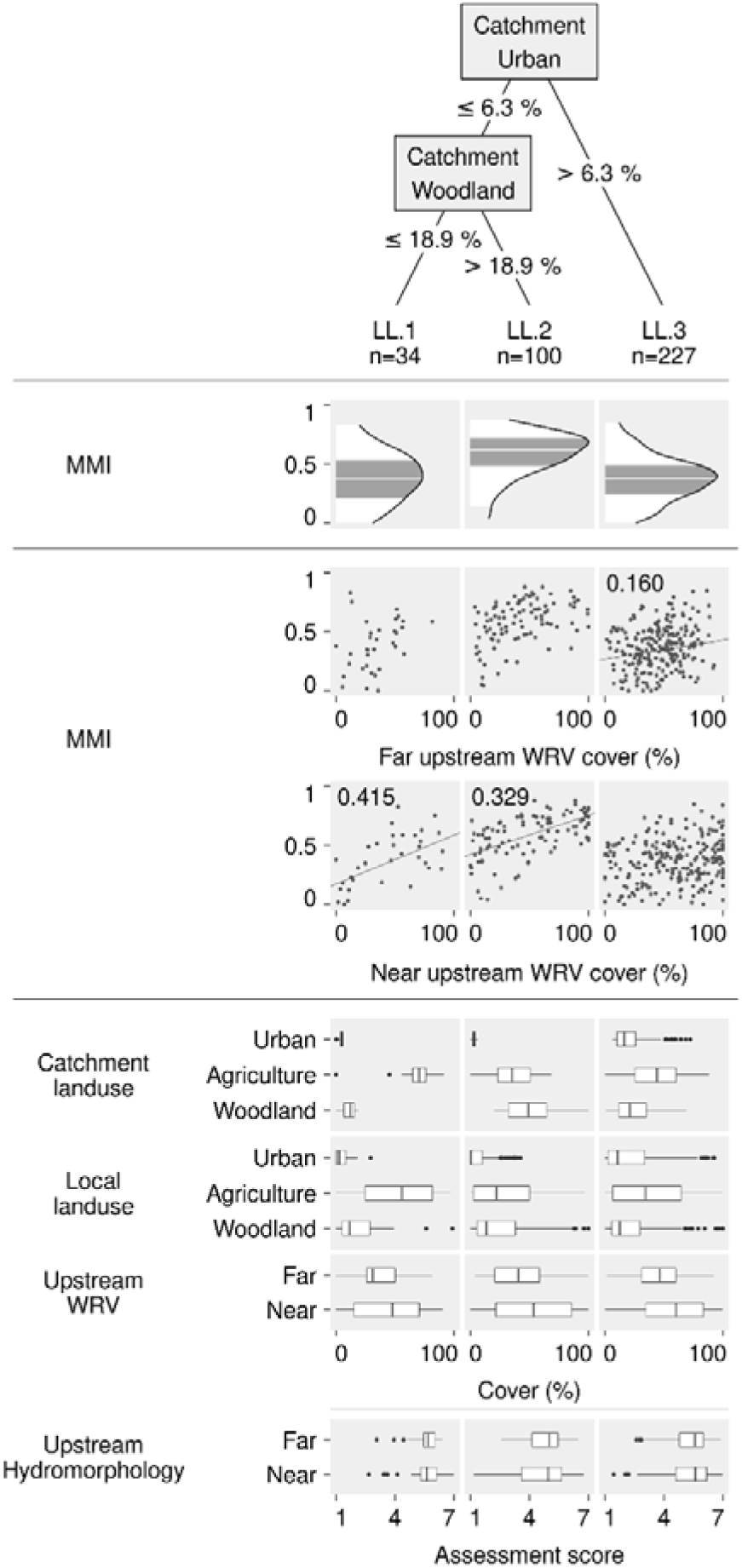
Partitioning tree for lowland sites. Density distributions of the macroinvertebrate multimetric index (MM) for each final subdataset (columns) with boxplot-like coloration of quantiles. Relationship between the MMI and the near and far upstream woody riparian vegetation (WRV) shown in scatterplots with significant effects indicated by regression coefficient and line. Distribution of all candidate partitioning variables given as boxplots.

In the lowlands, near upstream woody riparian vegetation (WRV) had the largest effect on macroinvertebrates’ ecological status in rural, agricultural catchments (n = 34; regression coefficient = 0.415). This subdataset LL.1 was characterized by low urban (≤ 6.3%; median = 4.9%; 75^th^-percentile = 5.7%) and low woodland cover (≤ 18.9%; median = 12.4%; 75^th^-percentile = 15.5%) in the catchment. Consequently, agriculture cover was high in the catchment (median = 70.3%, 25^th^-percentile = 65.1%) but also at the local scale (median = 55.9%; 75^th^-percentile = 81.9%).

Near upstream WRV had an intermediate effect on the MMI in rural, forested catchments (LL.2; n = 100; regression coefficient = 0.329) with low urban (≤ 6.3%; median = 3.1%; 75^th^-percentile = 4.0%) but much high woodland (> 18.9; median = 49.0%; 75^th^-percentile = 64.6%) cover in the catchment. Additionally, local woodland cover was slightly higher compared to subdataset LL. 1 (median = 13.5%; 75^th^-percentile = 38.0%).

WRV had the smallest effect on the MMI (LL.3; n = 227; regression coefficient far upstream = 0.160) in catchments with high urban cover (> 6.3%; median = 16.4%; 75^th^-percentile = 26.0%). Local urban cover was equally high, also compared to the other two subdatasets (median = 10.7%; 75^th^-percentile = 33.8%). Local woodland cover was similar to subdataset LL.1 (median = 12.6%; 75^th^-percentile = 29.9%) and woodland at the catchment scale was intermediate (median = 21.1%; 75^th^-percentile = 35.0%).

Since near upstream WRV was virtually un-correlated (and even negatively) with catchment woodland cover in subdataset LL.1 (Spearman’s *ρ* = −0.086; Table 1), the observed positive effect on the MMI was not simply due to sampling sites being located in forested areas. Conversely, near upstream WRV in LL.2 correlated moderately with catchment woodland cover (Spearman’s *ρ* = 0.404), implying that the smaller effect on the MMI might be partly due to positive effects of larger-scale forest cover. Finally, far upstream WRV in LL.3 correlated just weakly with catchment woodland (Spearman’s *p* = 0.291).

**Table 1:** Spearman’s *ρ* rank correlation coefficient between near and far upstream woody riparian vegetation (WRV) and catchment woodland cover for subdatasets with a significant effect (bold) from WRV on the macroinvertebrate multimetric index.

| Subdataset | Spearman's $\rho$ | |
| --- | --- | --- |
|  | near upstream WRV | far upstream WRV |
| LL.1 | **-0,086** | 0,115 |
| LL.2 | **0,404** *** | 0,547 *** |
| LL.3 | 0,128 | **0,291** *** |
| M.1 | 0,153 | **0,614** *** |
| M.2 | **-0,103** | 0,354 *** |
| M.5 | **-0,144** | 0,33 * |
| M.6 | 0,019 | **0,3** * |
| M.8 | 0,211 | **-0,066** |
| M.9 | -0,449 * | **-0,182** |
| M.10 | **0,044** | **0,24** * |

### 3.1 Mountain streams

Sites in mountain streams were split into eleven subdatasets (M.1 – M.11) by recursive partitioning based on four different partitioning variables (Fig. 4). Significant effects were found in seven of these subdatasets, with regression coefficients ranging from 0.149 to 0.995:

**Fig. 4:**
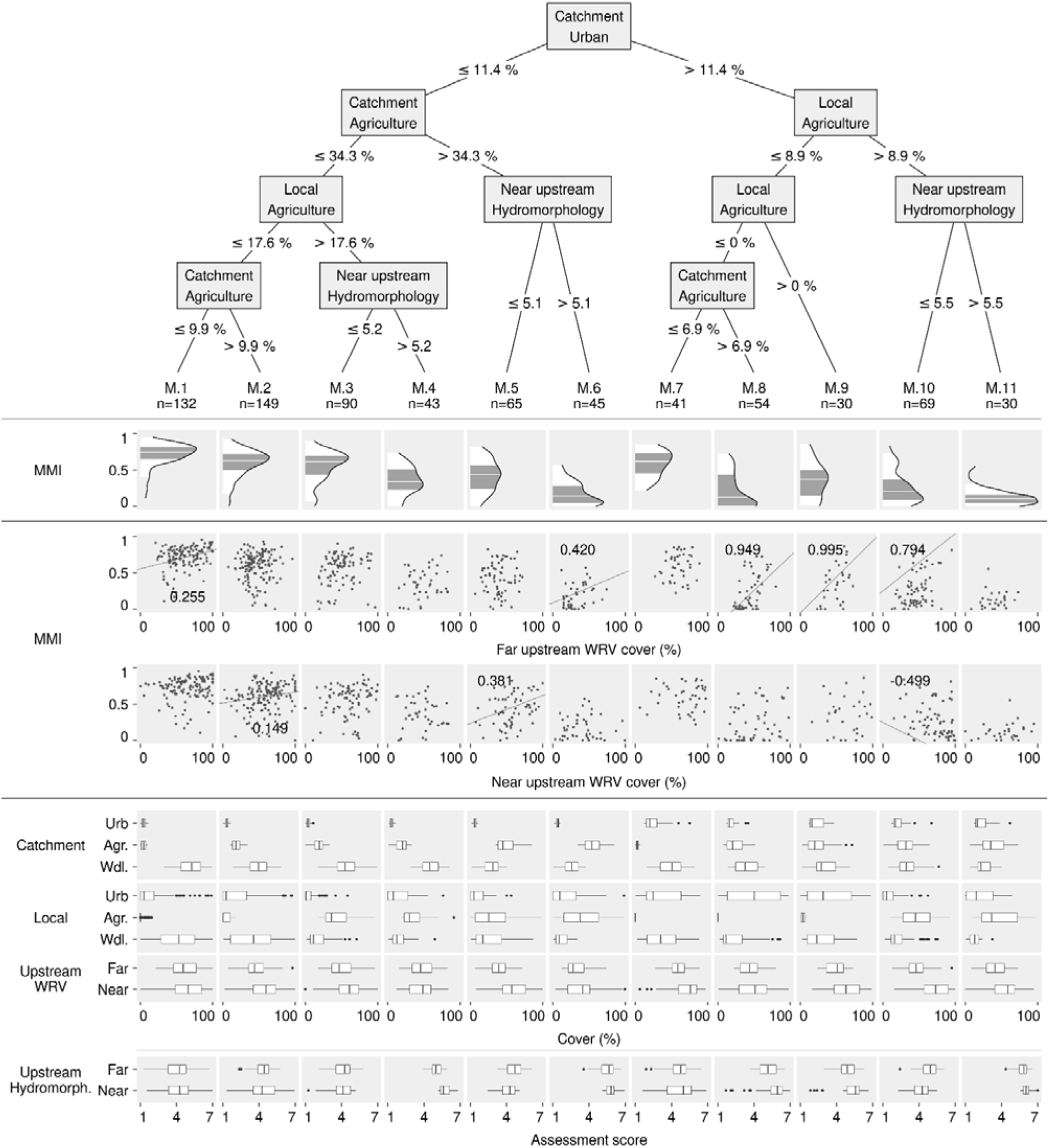
Partitioning tree for mountain sites. Density distributions of the macroinvertebrate multimetric index (MM) for each final subdataset (columns) with the boxplot-like coloration of quantiles. Relationship between the MMI and the near and far upstream woody riparian vegetation (WRV) shown in scatterplots with significant effects indicated by regression coefficient and line. Distribution of all candidate partitioning variables given as boxplots (Abbreviations: “Urb.” = urban, “Agr.” = agriculture, “Wdl.” = woodland).

Woody riparian vegetation (WRV) had a similarly large effect on the multimetric index (MMI) in two rural, agricultural subdatasets compared to lowland streams: Subdatasets M.5 (n = 65) and M.6 (n = 45) were characterized by low urban (≤ 11.4%; pooled median = 6.1% and 75^th^-percentile = 7.9%) and high agricultural cover (> 34.3%; pooled median = 49.2% and 75^th^-percentile = 61.2%) in the catchment. They were partitioned from each other using near upstream hydromorphology, which was substantially altered in M.5 (≤ 5.1; median = 4.2; 75^th^-percentile = 4.7), but even strongly degraded in M.6 (> 5.1; median = 5.8; 75^th^-percentile = 6.1). Besides these habitat conditions, the two subdatasets also differed in local woodland cover, which was intermediate in M.5 (median = 17.4%; 75^th^-percentile = 43.4%) and notably lower in M.6 (median = 8.7%; 75^th^-percentile = 20.1%). In subdataset M.5 with worse hydromorphology and higher local woodland cover, near upstream WRV had a positive effect on the MMI (regression coefficient 0.381). In subset M.6 with the less severe hydromorphological degradation and lower local woodland cover, far upstream WRV had a similar positive effect on the MMI (regression coefficient 0.420). Catchment woodland cover was moderate for both subdatasets (pooled median = 29.6%; 75^th^-percentile = 37.9%). WRV also had a significant but smaller effect on macroinvertebrates’ ecological status in rural, non-agricultural catchments, with a regression coefficients of 0.255 and 0.149 in subdatasets M1 and M2, respectively. These two subdatasets M.1 (n = 132) and M.2 (n = 149) were both characterized by low urban (≤ 11.4%; pooled median = 5.0% and 75^th^-percentile = 6.9%) and moderate agricultural cover in the catchment (≤ 34.3%) as well as very low agricultural cover locally (≤ 17.6%, pooled median = 0.0% and 75^th^-percentile = 5.0%). Consequently, local woodland cover was high in both subdatasets (M.1: median = 53.1%; M2: median = 42.6%).

Far upstream WRV had the largest effect on macroinvertebrates’ ecological status in urban catchments as observed in subdatasets M.8 (n = 54), M.9 (n = 30), and M.10 (n = 60). These three subdatasets were characterized by high catchment urbanisation (> 11.4%; pooled median = 16.2%; 75^th^-percentile = 23.6%). Sites in subdataset M.8 and M.9 where further characterized by low local agricultural cover (≤ 8.9) that was even completely lacking in M.8, which in turn featured intermediate catchment agriculture cover (> 6.9%; median ^= 20.3%; 75^th^-percentile = 33.9%). Rather high shares of local agriculture (> 8.9%; median = 45.1%; 75^th^-percentile = 65.2%) and less than strongly altered near upstream hydromorphology (≤5.5; median = 4.26; 75^th^-percentile = 4.72) characterized subdataset M.10. Far upstream WRV had exceptionally strong significant effects on the MMI in two of these subdatasets (regression coefficient M.8 = 0.949; M.9 = 0.995) where local urbanization was very high (pooled median = 42.9%; 75^th^-percentile=78.3%) while catchment (pooled median = 33.3%; 75^th^-percentile=54.5%) and local woodland (pooled median = 13.5%; 75^th^-percentile=39.5%) were intermediate at best. The third urban subdataset M.10, was the only one with significant effects from both near (regression coefficient −0.499) and far upstream (regression coefficient 0.794) WRV as well as the only with a negative effect (near upstream). Local (median = 16.2%; 75^th^-percentile = 26.6%) and catchment (median = 32.4%; 75^th^-percentile = 42.6%) woodland cover were both intermediate. Despite the highest amounts of catchment urbanisation of any subdataset with significant effects, local urbanisation was barely moderate (median = 5.1%; 75^th^-percentile = 14.4%).

In the two rural, agricultural subdatasets M.5 and M.6, the effects of near and upstream WRV were considered independent from large-scale woodland cover. Near upstream WRV was even negatively correlated with catchment woodland in M.5 (Spearman’s *ρ* = −0.114; Table 1), while far upstream WRV was positively but weakly related to catchment woodland in M.6 (Spearman’s *ρ* = −0.300).

In contrast, in the rural, non-agricultural subdataset M.1, far upstream WRV strongly correlated with catchment woodland (Spearman’s *ρ* = 0.614), indicating that the effects of WRV might be at least partly due to larger-scale forest cover. Yet in the second rural, non-agricultural subdataset (M.2), near upstream WRV correlated negatively with catchment woodland (Spearman’s *ρ* = −0.103).

In two of the urban subdatasets (M.8, M.9), far upstream WRV was also negatively correlated with catchment woodland cover, and only weakly positively correlated with catchment woodland cover in the third urban subdataset (M. 10).

## 4. Discussion

This study aimed to identify conditions, under which woody riparian vegetation (WRV) in a narrow buffer has the largest effects on the ecological status of macroinvertebrates (multimetric index, MMI). Despite using, to our knowledge the most detailed data on WRV in a large-scale study to date, there exist limitations to the approach. First, only percentage cover of certain landuse forms was assessed, which simplifies characteristics of more complex landscape elements (e.g. tree species, type of built-up area) and neglects temporal dynamics (e.g. forest development phase) as well as spatial arrangement. For instance, gaps in WRV are not accounted for and neither is it possible to perfectly distinguish WRV from wider forest cover by just comparing the narrow riparian corridor to local landuse and the entire catchment.

### 4.1 Rural landscapes

As hypothesized, WRV had a large positive effect on the ecological status of macroinvertebrates (multimetric index; MMI), in rural, agricultural landscapes. Results were consistent and regression coefficients similar in one subdataset in lowland (LL.1, regression coefficient 0.415) and two subdatasets in mountain streams (M.5 and M.6, regression coefficients 0.381 and 0.420). As the MMI ranges from 0 to 1 and is discretized evenly into five ecological status classes (high, good, moderate, poor or bad status), these coefficients imply that by managing woody riparian cover between 0 and 100%, without accompanying measures, the macroinvertebrate ecological status could be improved by as much as two status classes. This confirmed that woody riparian buffers are indeed a powerful tool for restoration in streams impacted by agricultural stressors. Given low catchment (median: 12.4 – 31.0%) and local (median: 8.6 – 17.4%) woodland cover and the lack of strong correlations between woodlands and WRV, the observed significant positive effects can be considered independent from larger-scale forest cover.

Other than expected, far upstream WRV was less important than near upstream WRV in agricultural landscapes. There was only one significant effect of far upstream WRV in one out of these three subdatasets (M.6). In the remaining two (LL.1, M.5) the significant positive effect was caused by near upstream WRV indicating that functions of WRV already were provided over a rather short distance of 500 m, which can substantially improve the ecological status of macroinvertebrate communities. This observation is in line with Kail et al. (2021), who observed that 400 m of shading by WRV results in a new thermal equilibrium of water temperature in lowlands. The sampling sites in subdataset M.6, where far upstream WRV had the significant effect, were highly morphologically degraded, suggesting that WRV at a larger spatial scale is necessary to compensate for instream habitat deficits.

In other rural but non-agricultural subdatasets, the effect of WRV on the ecological status was similar in lowland (LL.2, regression coefficient = 0.329) or somewhat lower in mountain streams (regression coefficient in M.1 = 0.255; M.2 = 0.149). Woodland cover in the catchment and locally around the sampling sites was much larger in these three subdatasets and correlated with WRV in two of them (LL.2, M.1). Therefore, the observed effects cannot be clearly attributed to WRV and might have been at least partly due to positive effects of large-scale forest cover. This would be consistent with other studies reporting strong positive effects of catchment woodland cover (Wahl et al., 2013).

### 4.2 Urbanized catchments

Other than expected, the effect of WRV on the ecological status was not clearly limited or superimposed by urban catchment landuse. Also, the percentage cover identified as the root split node well mirrored previously identified thresholds for urban cover with respect to the state of the macroinvertebrate community (Kail et al., 2012). Causes for the overall impact of urban areas are manifold (Walsh et al., 2005). For instance, increased runoff from impervious cover and flood prevention measures result in alterations to the hydrological regime. Furthermore, urban areas are sources for nutrients and hazardous substances, which eventually end up in streams. These impacts are evident in this study and reflected by generally lower multimetric (MMI) scores for subdatasets above the root split point (catchment urban cover) in both stream types.

However, within this limited range of low MMI scores, far upstream WRV still has a significant effect on the ecological status in urbanized catchments in lowland streams (LL.3). And even the by far largest effects are found in mountain streams in urbanized catchments (M.8, M.9, and M.10). Solely considering the regression coefficients one might expect that managing the woody riparian cover between 0 and 100% could improve the macroinvertebrate ecological status by as much as five status classes, i.e. from bad to high. This contradicts our expectations. However, given the lack of sites with high WRV cover and high ecological status such an extrapolation of the regression model over-interprets the results. Nevertheless, within the limited range of the data, results indicate that increasing WRV cover might be an appropriate restoration measure even in urban catchments. It seems that when there are virtually no adverse effects from agriculture to be buffered by near upstream WRV, the degree of naturalness in the far upstream riparian corridor, expressed by WRV cover, is a key determinant of macroinvertebrates’ ecological status at urban sites.

While the presence of far upstream WRV could be a proxy for the lack of near-stream urban pressures, the effect might also be due to functions provided by far upstream WRV, like decreasing water temperature or aiding aerial dispersal. These functions might – while not necessarily mitigating stressors related to urbanization like point source pollution and stormwater runoff – still improve habitat conditions rendering WRV a worthwhile restoration tool. In contrast to rural, agricultural settings, where near upstream WRV was most important, longer segments of WRV seem to be necessary to substantially improve habitat conditions in urban settings. Only in streams affected by urbanization in concert with agriculture and hydromorphological degradation even the positive effects of far upstream WRV are limited or superimposed by this multiple pressure situation (M.11).

Subdataset M.10 is furthermore special due to the, at first counterintuitive, negative effect from near upstream WRV along with the positive effect from far upstream WRV. Further inspection revealed a spatial cluster of sites within M.10 (Fig. 5), characterized by near upstream WRV upwards of 50% that nevertheless maintains poor MMI scores. This spatial cluster is located in the vicinity of Frankfurt am Main, a major metropolitan area, in small stream tributaries to the Nidda, which discharges to the river Main, as well as other close-by smaller direct tributaries to the Main. We suspect some local effect not accounted for in the data to be responsible. Excluding these sites from the subdataset, a positive effect would exist for both scales of woody riparian vegetation.

**Fig. 5:**
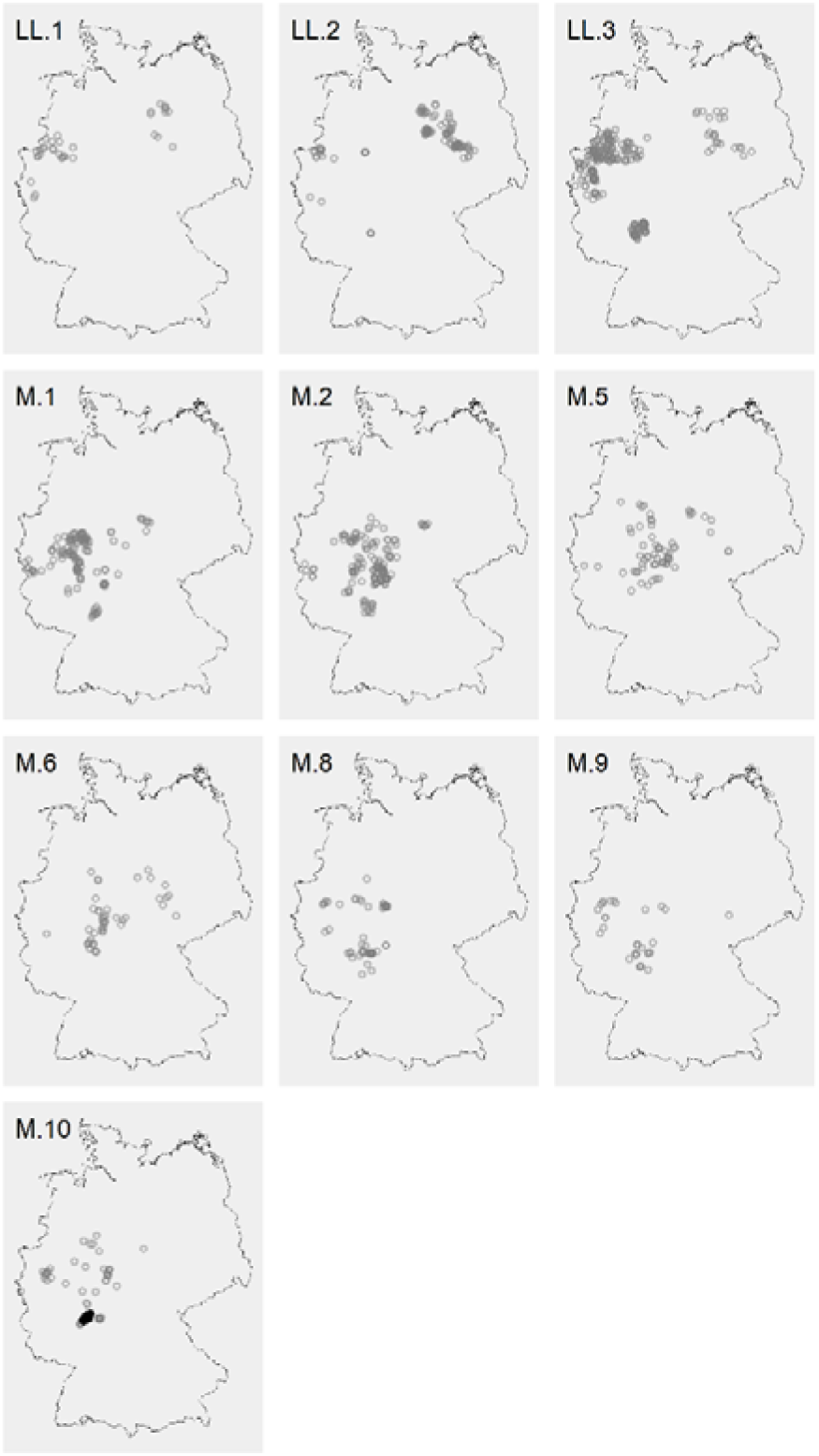
Sites in subdatasets with significant effects of woody riparian vegetation (WRV). Lowland sites in the top row. Sites in subdataset M.10 contributing to a negative effect of WRV in black.

## 5. Conclusion

Numerous small scale studies confirmed the beneficial effects of WRV on macroinvertebrates, while the results of recent large scale studies (Le Gall et al. 2021; Palt et al., 2022) question the effectiveness of woody riparian vegetation (WRV) to improve the ecological status. Our findings clearly reveal that effects of WRV on ecological status are large, but context specific as they differ in magnitude and scale according to catchment landuse, local landuse and hydromorphology. While the identification of context-specificity of the relationship between woody riparian vegetation and the macroinvertebrate community is hardly surprising, this analysis first succeeds in confirming underlying assumptions using a large dataset.

In streams mainly impacted by catchment urbanization, longer upstream reaches bordered by WRV seem to be necessary to substantially improve habitat conditions enhancing macroinvertebrates’ ecological status in the range from bad to moderate conditions. Establishing WRV is potentially particularly relevant for river management in rural, agricultural settings, where an increase in WRV from 0 to 100% can improve ecological status by up to two classes. Thus developing WRV can be an effective measure to reach good ecological status. We conclude that establishment of WRV is a key measure in the management and restoration of small streams, which is effective and easily applicable.

## 6. Authors’ contributions

MP, DH and JK conceived the ideas; MP designed the methodology and analysed the data; MP and JK led the writing of the manuscript. All authors contributed critically to the drafts and gave final approval for publication.

## Statement on inclusion

Our study relied on the assemblage of available data from the authors’ own country and therefore no data collection was needed. The research was conducted in cooperation with local state authorities and agency stakeholders were involved in the greater study project.

## 7. Acknowledgements

We thank the German Federal Ministry of Education and Research (01LC1618A) for funding the 2015-2016 BiodivERsA COFOUND project OSCAR during which the dataset was assembled and also the Department for the Environment, Agriculture and Geology of the State of Saxony for funding the analysis. Especially, we thank Bernd Spähnhoff.

